# Clade 2.3.4.4b H5N1 influenza virus and SARS-CoV-2 seroprevalence among owned and feral cats in Philadelphia and surrounding communities

**DOI:** 10.64898/2026.07.03.736283

**Authors:** Gabrielle Scher, Kathrine Maguire, Caitlin Duffy, Katie Mina, Clara Malekshahi, Stephen D. Cole, Laura R.H. Ahlers, Jacob N. Wohlstadter, George B. Sigal, Roderick B. Gagne, Louise H. Moncla, Scott E. Hensley

**Author notes:** Correspondence to Louise H. Moncla, and Scott E. Hensley.

## Abstract

Clade 2.3.4.4b H5NX influenza viruses have spread widely in birds since 2020. In addition to causing disease in birds, these viruses have infected a variety of mammals, including humans. Clade 2.3.4.4b H5N1 viruses are currently causing an outbreak among dairy cattle in the United States, and it is important to determine if other mammals have been exposed to H5NX viruses. Cats, specifically outdoor and feral cats, frequently predate wild birds. Recent studies have shown that cats living on dairy cattle farms can be infected with H5N1. Here, we completed serological studies to determine if owned and feral cats living in an urban environment in the United States have evidence of past H5N1 exposures. We used multianalyte bead-based assays to measure clade 2.3.4.4b hemagglutinin (HA) antibody levels in serum samples collected in July 2023 to June 2025 from 417 feral and 228 owned cats from the greater Philadelphia area. We also measured antibody levels against a panel of HAs from other human and non-human influenza viruses, and the receptor binding domain (RBD) of the severe acute respiratory syndrome coronavirus 2 (SARS-CoV-2). We completed additional H5N1 and SARS-CoV-2 neutralization assays using samples that had detectable antibodies in the multianalyte bead-based assays. One cat (0.16%) was positive for H5 antibodies and twenty cats (3.1%) were positive for SARS-CoV-2 antibodies in both binding and neutralization assays. These data suggest that cats in the Philadelphia area have not been routinely exposed to clade 2.3.4.4b H5N1 viruses but have been more commonly exposed to SARS-CoV-2.

**Importance:** Highly pathogenic H5NX avian influenza viruses have caused widespread disease and mortality in both wild and domestic birds, and have infected marine mammals, cattle, and humans. Recent studies have demonstrated that cats can also be infected with H5NX viruses, but it is unknown if these infections are common. In this report, we completed a seroprevalence study and found that H5NX exposure in feral and owned cats in the Philadelphia area appears to be rare. We also completed SARS-CoV-2 serological experiments and found that 3.1% of cats possessed antibodies reactive to this virus.

## INTRODUCTION

Highly pathogenic avian influenza viruses (HPAIVs) cause infections with high mortality rates in both wild and domestic bird populations^1,2^. The 1996 A/goose/Guangdong/1/96 (Gs/GD) lineage of H5 HPAIVs have become endemic in birds and periodically cause outbreaks^3,4^. While originally circulating in Asia, the Gs/GD lineage of H5 HPAIVs have spread via migratory birds across the globe^2,3^. Clade 2.3.4.4b H5NX viruses of the Gs/GD lineage have circulated widely since 2020 leading to high mortality rates in wild and domestic bird populations^5,6^. Additionally, clade 2.3.4.4b H5 HPAIVs have infected a variety of mammals, including mink, cattle, marine mammals, and humans^7–10^. Clade 2.3.4.4b H5 HPAIVs bind poorly to human upper airway cells^11,12^, but viral mutations from continued mammalian infection could lead to more efficient replication and increase transmission among humans^13–15^.

One factor that determines the risk of human infection with HPAIVs is their prevalence in animals with regular human contact. Cats are susceptible to avian influenza viruses both experimentally^16^ and naturally^17^. In 2016, a cat shelter in New York City experienced an outbreak of an H7N2 avian virus that infected around 500 cats and spread to a veterinarian at the shelter^18^. During the current H5N1 cattle outbreak, two cats living on an affected dairy farm were infected with clade 2.3.4.4b H5N1 virus, developed severe neurological symptoms, and succumbed to infection^19^. Additionally, two exclusively indoor cats whose owners who work on dairy farms suffered neurological symptoms and tested positive for H5N1 clade 2.3.4.4b viruses^20^. Further, a veterinarian had serological evidence of H5N1 infection following exposure to an infected cat suggesting that cat-to-human infections are possible^21^.

Serological studies have also demonstrated that some cats have serum antibodies that recognize avian influenza viruses. A 2020 serological study in the Netherlands found that 7.3% of shelter cats tested positive for H5 antibodies via H5 enzyme-linked immunosorbent assay (ELISA), and 1.6% were positive via nanoparticle based hemagglutination inhibition assay (HAI)^22^. In Spain, feral cats tested from March 2022-March 2023 in urban and peri-urban areas with confirmed wild bird HPAIV infections had a 2.19% positivity rate for H5 antibodies by ELISA for H5NX viruses^23^. A 2024 serological study in the Netherlands found that rural feral cats had an 11.8% positivity rate for H5 antibodies via ELISA alone and 9.3% positivity rate for H5 antibodies via both ELISA and HAI assays^24^.

Another concern with mammalian species that are susceptible to avian influenza is their ability to promote mammalian adaptation. It has been shown that cats express both alpha-2,3 and alpha-2,6 sialic acid-linked receptors in various tissues^25^, which could lead to the selection of H5N1 adaptive substitutions that enhance binding to human airway cells. A separate study of an H5N1 outbreak in cats in Poland in 2023 described that viruses isolated had mammalian adaptive mutations^26^. Thus, cats potentially represent a susceptible mammalian population with close contact to both humans and wild birds that could serve as an intermediary to spread HPAIVs to humans.

In this study, we examined clade 2.3.4.4b H5N1 seroprevalence using sera samples from owned and feral cats in the Philadelphia area. Although our study primarily focused on influenza viruses, cats are also susceptible to other widely circulating pathogens, including severe acute respiratory syndrome corona virus 2 (SARS-CoV-2)^27^. Thus, we also measured the seroprevalence of SARS-CoV-2 in these cats.

## RESULTS

### Sample collection and overview of serological approach

We collected serum samples from 315 feral and 228 owned cats in the greater Philadelphia area from July 2023 to May 2024. An additional 102 feral cat serum samples were collected from April 2025 to June 2025. We also obtained 11 naïve cat plasma samples from a specific pathogen free facility at Colorado State University in Fort Collins, CO. We first screened each sample using a multiplex bead-based assay to measure antibodies that bound to different viral antigens. We then completed additional neutralization assays with samples that had antibodies that bound to different viral antigens in the multiplex bead-based assays.

### Detection of binding antibodies in cat samples

We established multiplex bead-based assays that simultaneously measure antibody binding to HAs from a clade 2.3.4.4b H5N1 virus, a clade 1 H5N1 virus, and a low pathogenic avian influenza (LPAI) H5N1 virus. Within the same assays, we also measured antibody binding to HAs from influenza viruses that circulated seasonally in humans, a canine H3 from an H3N2 virus that circulates in dogs, a feline H7 from the 2016 shelter outbreak^18^, and the spike receptor binding domain (RBD) of two SARS-CoV-2 variants. As a positive control, we measured binding to the rabies virus glycoprotein (RABV-G) since the RABV vaccine is recommended for owned cats in the United States^28^. The antigens included in our multiplex bead-based antibody binding assay are summarized in **Table 1**.

**Table 1.** Luminex assay antigen panel.

| Antigen Abbreviation | Strain | Category of Antigen |
| --- | --- | --- |
| H5/2004 | A/Vietnam/1203/2004 | H5 Clade 1 virus |
| H5/2020 | A/Red Knot/Delaware Bay/404/2020 | H5 LPAIV |
| H5/2022 | A/Pheasant/New York/22-009066-001/2022 | H5 Clade 2.3.4.4b virus |
| H1/2022 | A/West Virginia/30/2022 | seasonal influenza virus |
| H3/2016 | A/Singapore/2016 | seasonal influenza virus |
| H3/2021 | A/Darwin/9/2021 | seasonal influenza virus |
| IBV/2017 | B/Colorado/06/2017 | seasonal influenza virus |
| IBV/2021 | B/Austria/1359417/2021 | seasonal influenza virus |
| H3/Canine/2023 | A/Canine/Minnesota/23-012684-001/2023 | additional influenza virus |
| H7/Feline/2016 | A/Feline/New York/WVDL-20/2016 | additional influenza virus |
| SARS-CoV-2 Wuhan | Wuhan-1 | SARS-CoV-2 |
| SARS-CoV-2 XBB.1.5 | XBB.1.5 | SARS-CoV-2 |
| Rabies G | Pasteur vaccins | Positive Control |

Antibodies in samples from cats living in a specific pathogen free facility bound poorly to each antigen (**Figure 1a**). Samples from owned (**Figure 1b**) and feral (**Figure 1c**) cats were considered to be seropositive to a specific antigen when the background subtracted mean fluorescence intensity (MFI) antibody binding values were ten times higher than the average background subtracted MFI values from naïve cats (**Table 2**). Using these criteria, we found that sera from 3.5% of owned (**Figure 1b**) and 2.64% of feral (**Figure 1c**) cats had antibodies that bound to at least one of the H5s tested. Surprisingly, sera from 60 (26.32%) of owned cats had antibodies that bound to HA from a human H1N1 virus, but very few samples (8, 3.51%) had antibodies that bound to HAs from the other human influenza viruses that we tested (**Figure 1b**). Seropositivity to at least one of the seasonal influenza viruses was comparable in owned cats (29.39%) and feral cats (25.18%) (**Figure 1b-c**). Seropositivity to the canine H3 and feline H7 were low overall, with only 3 (1.32%) owned cats and 6 (1.44%) feral cats testing positive for at least one of the canine or feline HAs (**Figure 1b-c**). Seropositivity to at least one SARS-CoV-2 RBD protein was similar in owned cats (12.23%) and feral cats (8.63%) (**Figure 1b-c**). As expected, a majority of the owned cats (94.74%) had antibodies that bound to RABV-G (**Figure 1b**, **Table 2**), while fewer feral cats (14.87%) had antibodies that bound to RABV-G (**Figure 1c**, **Table 2**).

**Figure 1.**
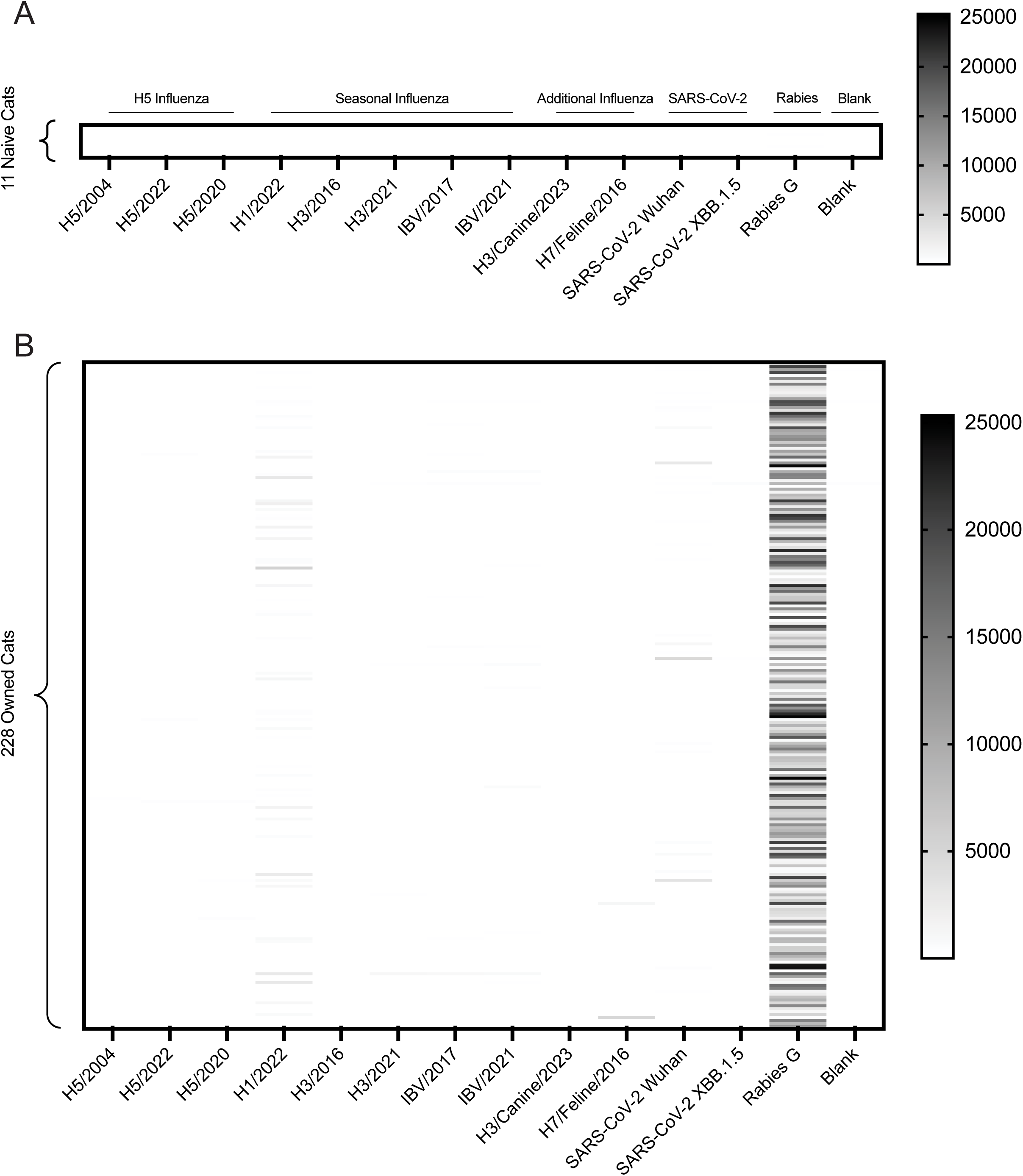
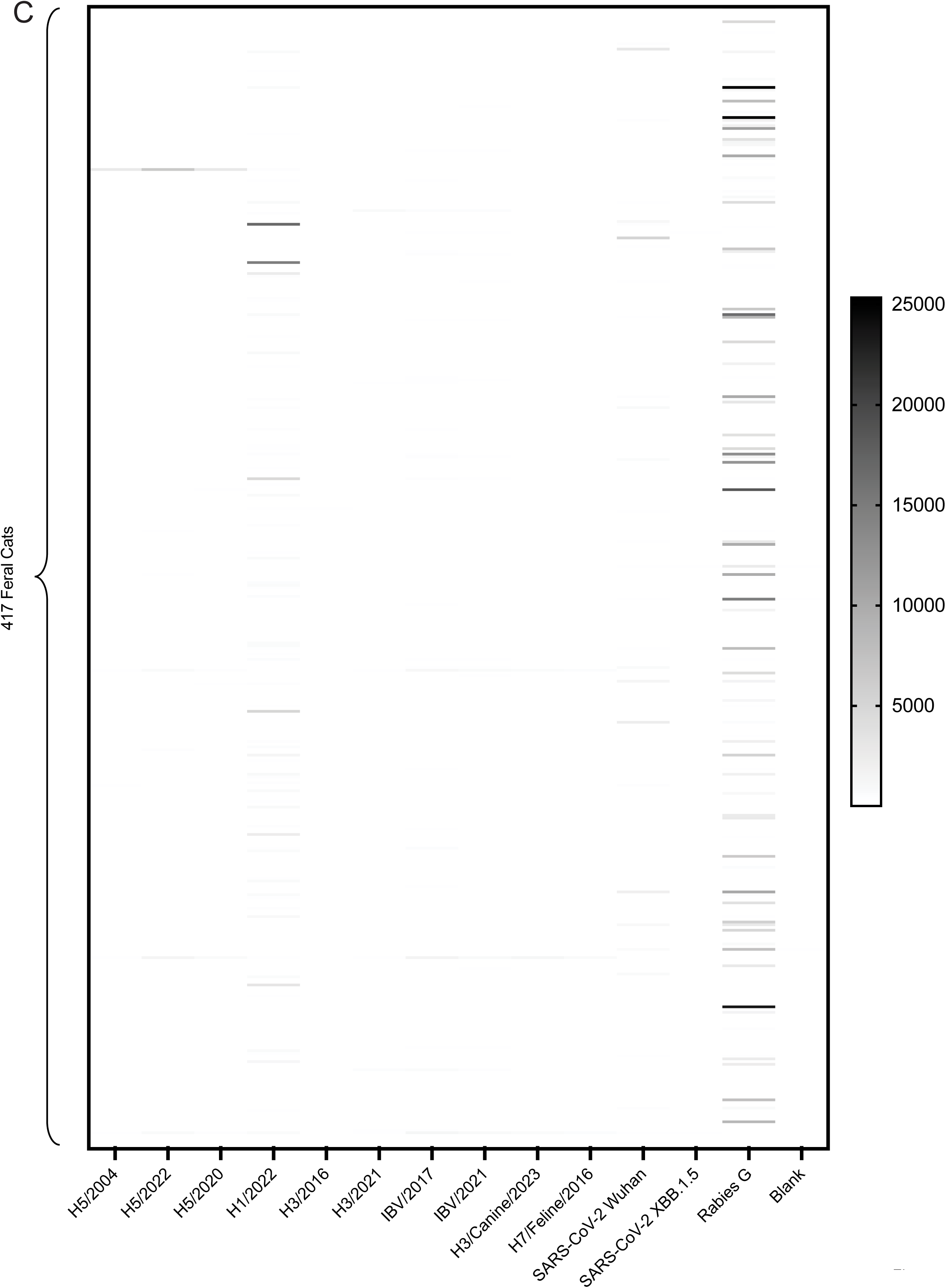
Owned and Feral Cats Have Binding Antibodies Against a Variety of Viral Antigens. Heat maps depicting the results of the Luminex assay for (**A**) naive, (**B**) owned, and (**C**) feral cat samples. The values in the heat map represent the background subtracted geometric mean of the mean fluorescent intensity (MFI). Samples were considered positive for an antigen if their MFI was 10 times higher than the average MFI for all the naïve samples for that antigen, unless that value was below 100, in which case an MFI of 100 was used as the cutoff for positivity. The legend for the heatmap represents the MFI values. Strain abbreviations are defined in **Table 1**.

**Table 2.** Luminex and neutralization positivity rates.

|  | Owned Cats | Feral Cats | Naïve Cats |
| --- | --- | --- | --- |
| **H5 Luminex +** | 7/228 (3.07%) | 7/417 (1.68%) | 0/11 (0%) |
| **H5 Luminex+/Neut +** | 0/228 (0%) | 1/417 (0.24%) | NT |
| **H1 Luminex +** | 60/228 (26.32%) | 94/417 (22.54%) | 0/11 (0%) |
| **H1 Luminex+/Neut +** | 4/226* (1.77%) | 19/417 (4.56%) | NT |
| **SARS-CoV-2 Luminex +** | 28/228 (12.28%) | 36/417 (8.63%) | 0/11 (0%) |
| **SARS-CoV-2 Luminex+/Neut +** | 10/228 (4.39%) | 10/417 (2.40%) | NT |
| **Rabies Luminex +** | 216/228 (94.74%) | 62/417 (14.87%) | 0/11 (0%) |
| **Rabies Luminex+/Neut +** | NT | NT | NT |

Neut – Neutralization; NT – Not tested
\* 2 samples that were H1 positive by Luminex could not be tested in neutralization assay because there was not enough sample.

### Identification of samples with neutralizing antibodies

Some influenza virus antibodies recognize epitopes that are conserved between different viral subtypes. For example, some antibodies elicited by H1N1 and H2N2 viruses can bind to the HA stalk domain of H5 viruses^29^. Consequently, our multiplex bead-based assays cannot distinguish between subtype-specific and cross-subtype antibodies. We therefore completed additional influenza virus neutralization assays that primarily detect antibodies recognizing strain-specific epitopes on the variable top of HA^30,31^. We focused on samples that were positive in our multiplex bead-based assay against the clade 2.3.4.4b H5 and the seasonal H1. We also completed SARS-CoV-2 neutralization assays using samples that had antibodies that bound to the SARS-CoV-2 Wuhan-1 spike RBD protein in the multiplex bead-based assays. As a control for the neutralization assays, we included samples that were negative in the multiplex bead-based assays.

Only 1 sample from feral cats (0.24%) and no samples from owned cats possessed clade 2.3.4.4b H5 neutralizing antibodies (**Figure 2a**, **Table 2**). This suggests that most of the antibodies that bound to clade 2.3.4.4b H5 in multiplex bead-based assays were not likely elicited by clade 2.3.4.4b H5 viruses, but rather other influenza virus subtypes or other H5 antigenic variants. Similarly, only 4 (1.78%) owned and 19 (4.56%) feral samples that bound to H1 viruses in the multiplex bead-based assays were able to neutralize H1N1 virus, with little correlation between binding and neutralizing titers (**Figure 2b**, **Table 2**). Of note, 2 of the owned cat samples that were positive for H1 binding could not be tested for neutralization due to low sample volumes. It is possible that the H1 binding antibodies that we identified were elicited by an H1N1 strain that is antigenically distinct compared to the H1N1 strain used in our neutralization assays. It is also possible that antibodies bound non-specifically to the H1N1-coated beads. Unlike what we observed for influenza viruses, many samples that had antibodies that bound to the SARS-CoV-2 spike protein also neutralized virus that was pseudotyped with the SARS-CoV-2 spike, with a strong correlation between binding and neutralizing antibodies (**Figure 2c**, **Table 2**).

**Figure 2.**
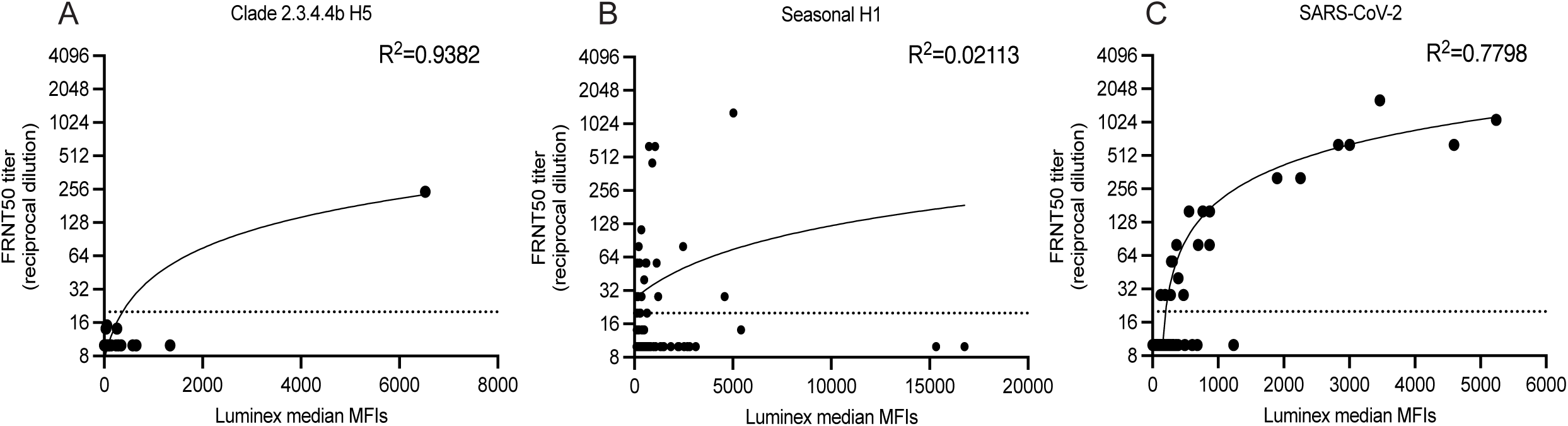
Luminex and Neutralization Assay Data Correlates Well for H5 and SARS-CoV-2. Luminex mean fluorescent intensity (MFI) values were plotted against 50% foci reduction neutralization test (FRNT50) titers (reciprocal serum dilution that inhibits 50% infection) for samples tested in neutralization assays for (**A**) clade 2.3.4.4.b H5 virus (H5 A/Pheasant/New York/22-009066-001/2022), (**B**) Seasonal H1 (H1 A/Wisconsin/50/2022), and (**C**) SARS-CoV-2 (SARS-CoV-2 Wuhan-1). Dotted line indicates the limit of detection for the neutralization assay. R^2^ values were determined by simple linear regression with a 95% confidence interval. P values were as follows: <0.0001 for H5 and SARS-CoV-2, and 0.0740 for H1.

To further validate our findings that there are low H5 positivity rates in cats in the Philadelphia area, we performed an H5 bridging serology assay. This assay measures antibodies from a sample that bind to both a capture antigen and detection antigen. We tested all samples that were 2.3.4.4b H5 positive in our multiplex bead-based assay (7 owned and 7 feral cats), and 66 negative samples, including a sample from a naïve laboratory cat (**Figure 3**). Samples were prepared both with and without an H1 blocker for testing. The H1 blocker in the assay adsorbs all H1-H5 cross-reactive antibodies. Most samples had low H5-reactivity in this assay with and without the H1 blocker (**Figure 3**). The single feral cat with H5 neutralizing antibodies had high levels of binding antibodies in this assay regardless of H1 blocker treatment (**Figure 3**).

**Figure 3.**
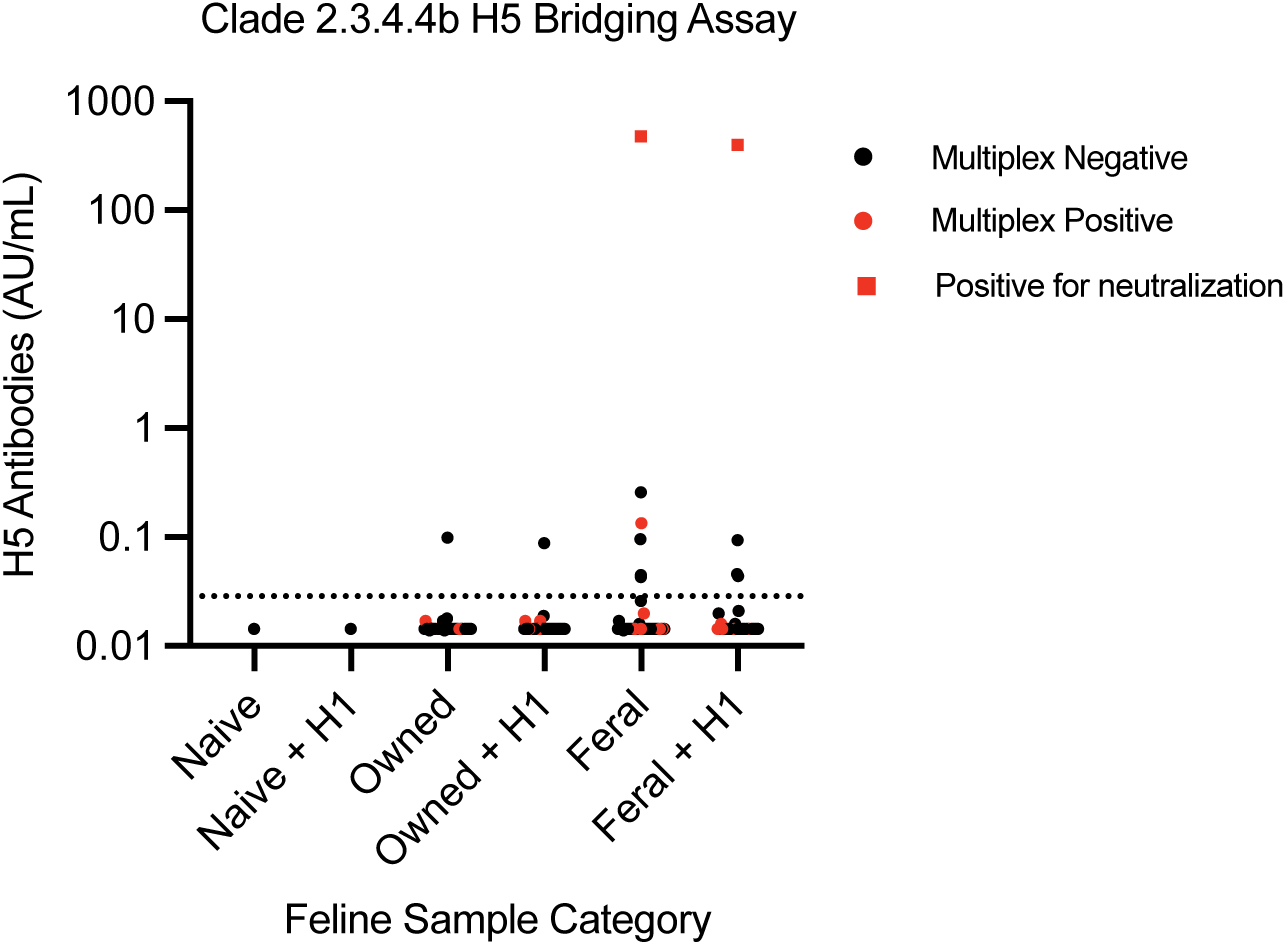
H5 Bridging Serology Assay Also Shows Low H5 Binding Antibody in Cats. The results from the H5 bridging serology assay for each sample are represented as arbitrary units (AU)/mL. Samples were considered positive when their AU/mL value is 2 times or above the limit of detection for the assay (0.01432 AU/mL). This cutoff is represented by the dotted line. Samples are separated by sample type (i.e. naïve, owned, or feral) and whether that sample condition received the H1 blocker (+ H1) or not. Individual samples are noted by circles, with the exception of samples that are neutralizing, which are indicated by squares. Samples that are negative for antibodies by the multiplex bead-based assay are in black, while samples that are positive for antibodies by the multiplex bead-based assay are in red.

Overall, these data suggest that most feral and owned cats in our study were not likely exposed to H5 HPAIV. While it was not our primary focus of this study, we found that over 2.4% of both feral and owned cats had antibodies that bound to and neutralized SARS-CoV-2 virus, suggesting that cats are more commonly exposed to this virus.

## DISCUSSION

In this study, we collected sera from feral and owned cats in the Philadelphia area and measured antibody binding to a panel of 13 viral antigens, completed additional neutralization assays with a subset of samples and performed a bridging serology assay against a 2.3.4.4b H5 to validate our results. Our primary goal was to determine if cats living in the Philadelphia area have been routinely exposed to clade 2.3.4.4b H5 HPAIVs. While we found that 3.07% of owned cats and 1.68% of feral cats possessed serum antibodies that could bind to clade 2.3.4.4b H5 in our multiplex bead-based assays, none of the samples from owned cats and only 0.24% of samples from feral cats (1 cat) had antibodies that neutralize a clade 2.3.4.4b virus. This suggests that the cats in our study were likely exposed to influenza viruses that have an HA that shares some conserved epitopes with clade 2.3.4.4b H5 viruses, but that infections with 2.3.4.4b H5 viruses in these cat populations were rare. Our findings are in agreement with other studies in cats that also found higher amounts of H5 binding antibodies compared to H5 neutralizing antibodies^22,24^.

Most cats sampled in this study reside in an urban environment. It is possible that cats living in urban environments are not exposed to avian influenza viruses as much as cats that are living on farms or more rural areas. Similar to our study, a recent report demonstrated low H5 seroprevalence rates in cats living in an urban area of Spain^23^. Consistent with the idea that urban and rural cats might have different rates of H5 exposures, a recent surveillance study from the Netherlands demonstrated that H5 positive cats were more likely to reside in rural areas such as nature reserves and farms^24^. Another study in France also found that outdoor cats residing in areas outside of cities had a relatively high H5 positivity rate (2.6%)^32^. Additional surveillance and serological studies should further investigate the association between urban and rural settings and risk of avian influenza virus infections of cats.

We do not fully understand why we observed that many cats in our study had serum antibodies that bound to H1 but could not neutralize H1N1 viruses. The simplest explanation is that these antibodies were elicited by an antigenically distinct H1N1 strain, and that we detected antibodies that recognized non-neutralizing conserved epitopes on the H1 used in our study.

Other studies have shown high H1 positivity rates in domestic cats for other H1 antigens, such as the H1 from the H1N1pdm2009 strain^22,24^. We cannot rule out the possibility that the high number of H1 positive samples is due to an artifact of antibodies binding non-specifically to H1 antigens coupled to beads. This phenomenon has been documented previously for cat samples tested in an enzyme-linked immunosorbent assay for feline immunodeficiency virus^33^. Many of the feral cats (62, 14.87%) in our study were positive for RABV-G antibodies. RABV is nearly 100% fatal in mammals once symptoms appear, and during infection the virus evades both the interferon and adaptive immune systems, preventing the production of antibodies^34,35^. Thus, the feral cats in our study that had RABV-G antibodies were likely vaccinated at some point before sampling. Consistent with this, many of the feral cats sampled in our study are part of cat colonies or shelters that vaccinate cats in their care, even if they are later released.

We found that 4.39% of owned and 2.4% of feral cats possessed both binding and neutralizing antibodies against SARS-CoV-2. Cats have been shown to be susceptible to SARS-CoV-2, experimentally^36^ and through human-to-cat transmission^37^. Our data are consistent with other studies that have found similar SARS-CoV-2 seroprevalence rates in cats^38–41^. There has only been one suspected case of cat-to-human SARS-CoV-2 transmission^42^. Together, these findings indicate that SARS-CoV-2 continues to infect animals, with cases in feral cats suggesting potential continued circulation in wildlife^43^.

Additional studies should be completed to continue monitoring H5N1 exposures of cats and other mammals that routinely interact with birds. Active surveillance studies, as well as serological studies can be used to carefully quantify the number of animals that are exposed to H5N1 viruses. This information can be used to better understand H5N1 host range and will be useful for informing pandemic risk assessments.

## Materials and Methods

### Sample Collection

Residual sera from owned cats were collected from Ryan Veterinary Hospital at the University of Pennsylvania. Cats presented at the hospital for unrelated care, and residual sera from existing blood draws was frozen and used in the study. For each cat, detailed patient data were collected including information on indoor/outdoor status, rabies vaccination status, breed, sex, and age.

Feral cat sera were collected as part of a trap, neuter, release program, from blood draws needed for testing of other feline pathogens and donated by the organization to this study. Sera for this study was refrigerated upon collection and stored at -80 within one week of collection.

### Recombinant protein production

To produce recombinant HA (rHA) proteins, the coding sequence of each HA was human codon optimized, the transmembrane domain was replaced with the Foldon T4 trimerization domain of T4 fibritin, an Avitag site-specific biotinylation sequence and hexahistidine tag^44^, and this sequence was synthesized into a pCMV-Sport6 vector (Twist bioscience). The following HA sequences were used: A/Singapore/INFIMH-16-0019/2016 (EPI780183), A/Darwin/9/2021 (EPI1883349), A/Vietnam/1203/2004 (EPI_ISL_10656749), B/Colorado/6/2017 (EPI1381168), B/Austria/1359417/2021 (EPI1845793), A/Pheasant/New York/22-009066-001/2022 (EPI ID 11971502), A/Red Knot/Delaware Bay/404/2020 (Genbank: MW875339.1), A/Canine/Minnesota/23-012684-001/2023 (Genbank: OQ990375.1), A/Feline/New York/WVDL-20/2016 (Genbank: MF978422.1), A/West Virginia/30/2022 (EPI2212467). A pCAGGS vector containing the receptor binding domain (RBD) of severe acute respiratory syndrome coronavirus 2 (SARS-CoV-2) wuhan-1 strain with a hexahistidine tag was a generous gift of Dr. Florian Krammer (Mt Sinai, NY). A pCMV-Sport6 containing the RBD of SARS-CoV-2 strain XBB.1.5 was produced as previously described^45^. These various plasmids were transfected in 293F cells using 293Fectin (Thermofisher) transfection reagent according to the manufacturer’s instructions. All recombinant HA plasmids were co-transfected with a plasmid encoding the neuraminidase (NA) from strain A/Puerto Rico/8/1934. Transfection supernatants were harvested after four days and proteins were purified using Ni-NTA agarose beads (Qiagen).

Rabies glycoprotein (rabies G), Pasteur vaccins strain (UniProt P08667) was purchased from Proteogenix to use as a positive control.

### Multiplex binding assay (Luminex Assay)

Recombinant proteins were first coupled to MagPlex beads (Luminex) at a ratio of 0.1nmol antigen per 1×10^6^ beads. The beads were mixed in blocking buffer (1X PBS, 0.1% tween20, 5% milk), added to the wells of the black, clear-bottom 96-well assay plates for a final concentration of 2500 beads of each bead per well in 50 µL and blocked overnight at 4°C. Heat inactivated sera were diluted to 1:400 in primary blocking buffer (1X PBS, 5% goat serum, 1% milk, 0.08% polyvinylpyrrolidone and 0.05% polyvinyl alcohol). The blocking buffer was removed from the beads, 50 µL of the diluted sera was added to the beads and incubated on a plate shaker for 1 h at 600 rpm. After incubation, plates were washed twice with PBS-TBN (1X PBS, 0.1% bovine serum albumin, 0.02% tween20, and 0.05% sodium azide) and biotin conjugated goat anti-cat IgG (H&L) (Rockland Immunochemicals) was added to each well at a concentration of 4 µg/mL in 50 µL. Plates were incubated for 30 min on the plate shaker and then washed twice with PBS-TBN. R-phycoerythrin (PE) conjugated streptavidin (BD Biosciences) was diluted 1:2000 in PBS-TBN, 50 µL was added to each well and the plate was incubated for 30 min on the plate shaker. After incubation, the plates were washed twice with PBS-TBN and 100 µL of PBS-TBN were added to each well. An Intelliflex (Luminex) was used to read the plates and the background subtracted geometric mean of the mean fluorescent intensities (MFI) from two independent replicates are reported.

### H5 *in vitro* neutralization assays

H5 neutralization assays were performed using replication deficient influenza viruses with the PB1 coding sequence replaced by GFP, where virus replication is dependent on cells expressing PB1, as previously described^46^. Viruses were recovered by transfecting a coculture of 293T-CMV-PB1 and MDCK-SIAT1-TMPRSS2-PB1 cells with bidirectional reverse genetics plasmids encoding PB2, PA, NP, M and NS from A/Puerto Rico/8/1934, HA segment from A/Pheasant/New York/22-009066-001/2022, NA segment from A/Astrakhan/3202/2020, GFP in the open reading frame of the PB1 gene and a plasmid encoding TMPRSS2. The multibasic cleavage site KRRKR in HA was replaced with a monobasic R as an added safety measure.

Media was changed at 20 h post-transfection to neutralization assay media (NAM; Medium 199 (Thermo Fisher), 0.01% heat-inactivated fetal bovine serum, 0.3% bovine serum albumin, 100 µg penicillin/mL, 100 µg streptomycin/mL, 100 µg calcium chloride/mL and 25 mM HEPES). Supernatants were harvested at 72 h post-transfection, clarified by centrifugation and used to expand virus on MDCK-SIAT1-TMPRSS2-PB1 cells. Serum was treated with receptor-destroying enzyme (RDE; Denka Seiken) and then heat inactivated. In a 96-well flat-bottom plate, serum was serially diluted two-fold with NAM. Virus was diluted to a previously determined concentration to produce an MFI around 1000 and added to the plates at a volume equal to that of the serum dilution. The serum-virus mixes were incubated at 37 °C in 5% CO_2_ for 1 h, after which 2.5 x 10^4^ MDCK-SIAT1-TMPRSS2-PB1 cells were added to each well. Cells were incubated with the serum-virus mixes for 42 h and then fixed with 4% paraformaldehyde.

An EnVision microplate reader was used to measure the GFP fluorescence intensities at an excitation wavelength of 485 nm and emission wavelength of 530 nm. Neutralization titers are reported as the highest reciprocal serum dilution that elicited a reduction of 50% or more in GFP levels compared to the virus only control^47^. Serum samples that did not reach a 50% reduction in GFP levels even at the highest dilution of 1:20 were reported as having a 50% neutralizing titer of 10.

### H1 *in vitro* neutralization assays

A/Wisconsin/50/2022 H1N1 virus was used for the H1 neutralization assays as it has the same HA amino acid sequence as A/West Virginia/30/2022. Virus was recovered by transfecting a coculture of 293T and MDCK-SIAT1 cells with bidirectional reverse genetics plasmids encoding PB1, PB2, PA, NP, M and NS from A/Puerto Rico/8/1934, and the HA and NA segments from A/Wisconsin/50/2022 (EPI_ISL_15724390). Media was changed to NAM at 20 h post-transfection and then supernatants were collected and clarified by centrifugation at 72 h post-transfection. Transfection supernatant was used to expand virus stock on MDCK-SIAT1 cells. As described previously, this virus was used in a foci reduction neutralization assay (FRNT)^48^.

Briefly, MDCK-SIAT1 cells were seeded in a flat-bottom 96-well plate at 2.5×10^4^ cells/well and incubated at 37 °C overnight. Serum samples were treated with RDE and heat-inactivated before being serially diluted in serum-free MEM (SF-MEM). Virus was diluted to a previously determined amount to produce 300 focus-forming units per well, added to the serum in an equal volume and the resulting serum-virus mix was incubated for 1 h at 37 °C. The previously seeded MDCK-SIAT1 cells were washed twice with SF-MEM, then overlaid with the serum-virus mix and incubated for 1 h at 37 °C. Cells were then washed once more with SF-MEM, overlayed with 50% SF-MEM, 1.25% avicel, 5 mM HEPES, and 50 µg/mL Gentamycin-Sulfate and incubated for 18 h at 37 °C. Following incubation, plates were fixed with 4% paraformaldehyde and then permeabilized with 0.5% Triton-X 100 in 1X PBS. Next, 5% milk in 1X PBS was used to block the plates for 1 h. Mouse anti-influenza A NP antibody (IC5-1B7, produced in-house) was diluted 1:20,000 in 5% milk, added to the plates and incubated for 1 h. Next, Rat anti-mouse HRP antibody (Southern Biotech: 1170-05) was diluted 1:30,000 in 5% milk, added to the plates and incubated for 1 h. Finally, TrueBlue TMB substrate was added to the plates for development and the plates were read on an ELISpot reader. Neutralization titers are reported as the highest reciprocal serum dilution that elicited a reduction of 50% or more in the number of foci compared to the virus only control. Serum samples that did not reach a 50% reduction in number of foci even at the highest dilution of 1:20 were reported as having a 50% neutralizing titer of 10.

### SARS-CoV-2 pseudotype *in vitro* neutralization assays

SARS-CoV-2 neutralization assays were performed using vesicular stomatitis virus (VSV) pseudotyped with SARS-CoV-2 spike protein. This virus was produced, and the assay performed as previously described^45,49,50^. In brief, 3.5×10^6^ 293T cells were seeded in collagen-coated 10cm dishes and incubated for 24 h at 37 °C. Cells were then transfected with 25 µg of a human codon optimized SARS-CoV-2 spike protein from the Wuhan-1 strain with an 18-residue truncation in the cytoplasmic tail. 24 h post-transfection, cells were infected with VSVΔG-VSV-G-RFP at an MOI of ∼2-4, incubated for 2-4 h and then the virus containing media was replaced with fresh media. 24-28 h post-infection, VSVΔG-SARS-COV-2-S-RFP pseudovirus was harvested and clarified by centrifugation. 2.5×10^4^ VeroE6-TMPRSS2 cells were seeded in collagen-coated 96-well plates and incubated at 37 °C overnight. Serum was heat inactivated and serially diluted two-fold in assay medium (DMEM with 10% FBS and 600ng/mL of anti-VSV-G antibody IE9F9). Virus was diluted in assay medium to a dilution previously determined to produce 200-300 foci/well and added to the serum in a volume equivalent to that of the serum. After a 1 h incubation at 37 °C, the serum-virus mix was overlaid on the VeroE6-TMPRSS2 cells and incubated at 37 °C for 21-22 h. Plates were then washed once with 1X DPBS and fixed with 4% paraformaldehyde. An S6 FluoroSpot Analyzer (CTL) was used to read and count RFP foci.

Neutralization titers are reported as the highest reciprocal serum dilution that elicited a reduction of 50% or more in the number of foci compared to the virus only control. Serum samples that did not reach a 50% reduction in number of foci even at the highest dilution of 1:20 were reported as having a 50% neutralizing titer of 10.

### H5 bridging serology assay

The H5 bridging serology assay was performed using a commercial kit following the manufacturer’s instructions as previously described (Meso Scale Discovery (MSD), Cat # K150AXJU)^51^. Briefly, a master mix was prepared containing equal parts of biotin-labeled H5 antigen (capture antigen) and SULFO-TAG™ labeled H5 antigen (detection antigen). Samples were diluted 1:10 with or without H1 blocker (MSD, Cat # R93BL) and then 40µL of sample was mixed with 40µL of master mix. After 1 h of incubation on a shaker, 50µL of the sample/master mix mixture was added to a pre-washed MSD GOLD 96-Well Small Spot Streptavidin SECTOR™ Plate. Following another 1 h shaking incubation, the plate was washed, 150µL of MSD GOLD™ Read Buffer B was added to the plate and electrochemiluminescence (ECL) was measured using an MSD MESO SECTOR^®^ S 600MM plate reader. Samples were quantified using a 7-point calibration curve of the kit’s high antibody control and a four-parameter-logistic (4PL) fit with 1/Y^2^ weighting. The limit of detection (LOD) for the assay was the concentration of the calibration curve that gave a signal 2.5 standard deviations above the blank sample signal, specifically 0.01432 AU/mL. Samples below the LOD are reported as the LOD value and reported concentrations were corrected for sample dilution. Samples were considered positive when their values were at least 2-fold above the LOD.

### Statistical Analysis

The correlations between the Luminex median MFIs and neutralization FRNT50 titers were determined by Simple Linear Regression with a 95% confidence interval. All statistics were performed in GraphPad Prism 10 software.

## Acknowledgments

This project has been funded in part with Federal funds from the National Institute of Allergy and Infectious Diseases, National Institutes of Health, Department of Health and Human Services, under Contract No. 75N93021C00015 (S.E.H. and L.H.M.) and in part by NIAID Award No. 1UH2AI176136 (L.R.H.A and G.B.S.).The authors would like to acknowledge Susan VandeWoude and Mary Nehring from Colorado State University who provided the specific pathogen free cat plasma samples used in this study.

## References

1 Swayne, D. & Suarez, D. Highly pathogenic avian influenza. Revue scientifique et technique-office international des epizooties 19, 463–475 (2000). 10.20506/rst.19.2.1230

2. Swayne, D., Suarez, D. & Sims, L. in Diseases of poultry (eds DE Swayne et al.) 210–256 (Wiley, 2020).

3 Röhm, C., Horimoto, T., Kawaoka, Y., Süss, J. & Webster, R. G. Do Hemagglutinin Genes of Highly Pathogenic Avian influenza Viruses Constitute Unique Phylogenetic Lineages? Virology 209, 664–670 (1995). 10.1006/viro.1995.1301

4 Lee, D. H., Criado, M. F. & Swayne, D. E. Pathobiological Origins and Evolutionary History of Highly Pathogenic Avian Influenza Viruses. Cold Spring Harb Perspect Med 11 (2021). 10.1101/cshperspect.a038679

5. European Food Safety Authority, E. C. f. D. P., et al. Avian influenza overview September – December 2022. EFSA Journal 21, e07786 (2023). 10.2903/j.efsa.2023.7786

6 Youk, S. et al. H5N1 highly pathogenic avian influenza clade 2.3.4.4b in wild and domestic birds: Introductions into the United States and reassortments, December 2021–April 2022. Virology 587, 109860 (2023). 10.1016/j.virol.2023.109860

7 Agüero, M. et al. Highly pathogenic avian influenza A(H5N1) virus infection in farmed minks, Spain, October 2022. Euro Surveill 28 (2023). 10.2807/1560-7917.Es.2023.28.3.2300001

8 Burki, T. Avian influenza in cattle in the USA. The Lancet Infectious Diseases 24, e424–e425 (2024). 10.1016/S1473-3099(24)00369-4

9 Tomás, G. et al. Highly pathogenic avian influenza H5N1 virus infections in pinnipeds and seabirds in Uruguay: Implications for bird–mammal transmission in South America. Virus Evolution 10, veae031 (2024). 10.1093/ve/veae031

10 Venkatesan, P. Human cases of avian influenza A(H5) in the USA. The Lancet Microbe, 100978 (2024). 10.1016/j.lanmic.2024.100978

11 Santos, J. J. S. et al. Bovine H5N1 influenza virus binds poorly to human-type sialic acid receptors. bioRxiv (2024). 10.1101/2024.08.01.606177

12 Chopra, P. et al. Receptor Binding Specificity of a Bovine A(H5N1) Influenza Virus. bioRxiv (2024). 10.1101/2024.07.30.605893

13 Moratorio, G. et al. H5N1 influenza: Urgent questions and directions. Cell 187, 4546–4548 (2024). 10.1016/j.cell.2024.07.024

14 Alkie, T. N. et al. Characterization of neurotropic HPAI H5N1 viruses with novel genome constellations and mammalian adaptive mutations in free-living mesocarnivores in Canada. Emerg Microbes Infect 12, 2186608 (2023). 10.1080/22221751.2023.2186608

15 Eisfeld, A. J. et al. Pathogenicity and transmissibility of bovine H5N1 influenza virus. Nature 633, 426–432 (2024). 10.1038/s41586-024-07766-6

16 Kuiken, T. et al. Avian H5N1 Influenza in Cats. Science 306, 241–241 (2004). 10.1126/science.1102287

17 Coleman, K. K. & Bemis, I. G. Avian Influenza Virus Infections in Felines: A Systematic Review of Two Decades of Literature. medRxiv, 2024.2004.2030.24306585 (2024). 10.1101/2024.04.30.24306585

18 Hatta, M. et al. Characterization of a Feline Influenza A(H7N2) Virus. Emerg Infect Dis 24, 75–86 (2018). 10.3201/eid2401.171240

19 Burrough, E. R. et al. Highly pathogenic avian influenza A (H5N1) clade 2.3. 4.4 b virus infection in domestic dairy cattle and cats, United States, 2024. Emerging infectious diseases 30, 1335–1343 (2024).

20 Naraharisetti, R. et al. Highly Pathogenic Avian Influenza A(H5N1) Virus Infection of Indoor Domestic Cats Within Dairy Industry Worker Households — Michigan, May 2024. MMWR Morb Mortal Wkly Rep 74, 61–65 (2025). 10.15585/mmwr.mm7405a2

21 Vaughan, A. et al. Serologic Evidence of Highly Pathogenic Avian Influenza A(H5N1) Virus Infection in a Veterinary Professional Exposed to an Infected Domestic Cat — Los Angeles County, California, December 2024–January 2025. MMWR Morb Mortal Wkly Rep 75, 215–220 (2026). 10.15585/mmwr.mm7517a1.

22 Zhao, S. et al. Serological Screening of Influenza A Virus Antibodies in Cats and Dogs Indicates Frequent Infection with Different Subtypes. Journal of Clinical Microbiology 58, 10.1128/jcm.01689–01620 (2020). 10.1128/jcm.01689-20

23 Villanueva-Saz, S. et al. Serological exposure to influenza A in cats from an area with wild birds positive for avian influenza. Zoonoses and Public Health 71, 324–330 (2024). 10.1111/zph.13085

24 Duijvestijn, M. B. H. M. et al. Highly pathogenic avian influenza (HPAI) H5 virus exposure in domestic cats and rural stray cats, the Netherlands, October 2020 to June 2023. Eurosurveillance 29, 2400326 (2024). 10.2807/1560-7917.ES.2024.29.44.2400326

25 Chothe, S. K. et al. Marked neurotropism and potential adaptation of H5N1 clade 2.3.4.4.b virus in naturally infected domestic cats. Emerg Microbes Infect 14, 2440498 (2025). 10.1080/22221751.2024.2440498

26 Domańska-Blicharz, K. et al. Outbreak of highly pathogenic avian influenza A(H5N1) clade 2.3.4.4b virus in cats, Poland, June to July 2023. Euro Surveill 28 (2023). 10.2807/1560-7917.Es.2023.28.31.2300366

27 Doliff, R. & Martens, P. Cats and SARS-CoV-2: A Scoping Review. Animals (Basel) 12 (2022). 10.3390/ani12111413

28 Squires, R. A., Crawford, C., Marcondes, M. & Whitley, N. 2024 guidelines for the vaccination of dogs and cats – compiled by the Vaccination Guidelines Group (VGG) of the World Small Animal Veterinary Association (WSAVA). Journal of Small Animal Practice 65, 277–316 (2024). 10.1111/jsap.13718

29 Krammer, F. & Palese, P. Influenza virus hemagglutinin stalk-based antibodies and vaccines. Curr Opin Virol 3, 521–530 (2013). 10.1016/j.coviro.2013.07.007

30 Temoltzin-Palacios, F. & Thomas, D. B. Modulation of immunodominant sites in influenza hemagglutinin compromise antigenic variation and select receptor-binding variant viruses. J Exp Med 179, 1719–1724 (1994). 10.1084/jem.179.5.1719

31 Smith, C. A., Barnett, B. C., Thomas, D. B. & Temoltzin-Palacios, F. Structural assignment of novel and immunodominant antigenic sites in the neutralizing antibody response of CBA/Ca mice to influenza hemagglutinin. J Exp Med 173, 953–959 (1991). 10.1084/jem.173.4.953

32 Bessière, P. et al. Cats as sentinels of mammal exposure to H5Nx avian influenza viruses: a seroprevalence study, France, December 2023 to January 2025. Euro Surveill 30 (2025). 10.2807/1560-7917.Es.2025.30.12.2500189

33 Moskaluk, A., Nehring, M. & VandeWoude, S. Serum Samples from Co-Infected and Domestic Cat Field Isolates Nonspecifically Bind FIV and Other Antigens in Enzyme-Linked Immunosorbent Assays. Pathogens 10, 665 (2021).

34 Davis, B. M., Rall, G. F. & Schnell, M. J. Everything You Always Wanted to Know About Rabies Virus (But Were Afraid to Ask). Annu Rev Virol 2, 451–471 (2015). 10.1146/annurev-virology-100114-055157

35. Rieder, M. & Conzelmann, K.-K. in Advances in Virus Research Vol. 79 (ed Alan C. Jackson) 91–114 (Academic Press, 2011).

36 Park, E. S. et al. The comparison of pathogenicity among SARS-CoV-2 variants in domestic cats. Sci Rep 14, 21815 (2024). 10.1038/s41598-024-71791-8

37 Hosie, M. J. et al. Anthropogenic Infection of Cats during the 2020 COVID-19 Pandemic. Viruses 13 (2021). 10.3390/v13020185

38 Selyemová, D. et al. Cats as a sentinel species for human infectious diseases - toxoplasmosis, trichinellosis, and COVID-19. Curr Res Parasitol Vector Borne Dis 6, 100196 (2024). 10.1016/j.crpvbd.2024.100196

39 Suwanpakdee, S. et al. Sero-epidemiological investigation and cross-neutralization activity against SARS-CoV-2 variants in cats and dogs, Thailand. Front Vet Sci 11, 1329656 (2024). 10.3389/fvets.2024.1329656

40 Thongyuan, S. et al. Seroprevalence of Anti-SARS-CoV-2 Antibodies in Cats during Five Waves of COVID-19 Epidemic in Thailand and Correlation with Human Outbreaks. Animals (Basel) 14 (2024). 10.3390/ani14050761

41 Fritz, M. et al. A Large-Scale Serological Survey in Pets From October 2020 Through June 2021 in France Shows Significantly Higher Exposure to SARS-CoV-2 in Cats Compared to Dogs. Zoonoses Public Health 72, 184–193 (2025). 10.1111/zph.13198

42 Sila, T. et al. Suspected Cat-to-Human Transmission of SARS-CoV-2, Thailand, July-September 2021. Emerg Infect Dis 28, 1485–1488 (2022). 10.3201/eid2807.212605

43 Goldberg, A. R. et al. Widespread exposure to SARS-CoV-2 in wildlife communities. Nat Commun 15, 6940 (2024). 10.1038/s41467-024-51220-0

44 Whittle, J. R. et al. Flow cytometry reveals that H5N1 vaccination elicits cross-reactive stem-directed antibodies from multiple Ig heavy-chain lineages. J Virol 88, 4047–4057 (2014). 10.1128/jvi.03422-13

45 Johnston, T. S. et al. Immunological imprinting shapes the specificity of human antibody responses against SARS-CoV-2 variants. Immunity 57, 912–925.e914 (2024). 10.1016/j.immuni.2024.02.017

46 Doud, M. B., Hensley, S. E. & Bloom, J. D. Complete mapping of viral escape from neutralizing antibodies. PLOS Pathogens 13, e1006271 (2017). 10.1371/journal.ppat.1006271

47 Arevalo, C. P. et al. A multivalent nucleoside-modified mRNA vaccine against all known influenza virus subtypes. Science 378, 899–904 (2022). 10.1126/science.abm0271

48 Gouma, S. et al. Middle-aged individuals may be in a perpetual state of H3N2 influenza virus susceptibility. Nature Communications 11, 4566 (2020). 10.1038/s41467-020-18465-x

49 Goel, R. R. et al. mRNA vaccines induce durable immune memory to SARS-CoV-2 and variants of concern. Science 374, abm0829 (2021). 10.1126/science.abm0829

50 Anderson, E. M. et al. Seasonal human coronavirus antibodies are boosted upon SARS-CoV-2 infection but not associated with protection. Cell 184, 1858–1864.e1810 (2021). 10.1016/j.cell.2021.02.010

51 Marquez, A. C. et al. Detection of Avian Influenza H5–Specific Antibodies by Chemiluminescent Assays. Emerg Infect Dis 32, 129–132 (2026). 10.3201/eid3201.251117

